# Inferring Models with Alternative Stable States from Independent Observations

**DOI:** 10.1101/2020.02.07.939413

**Authors:** Edward W. Tekwa, Martin Krkošek, Malin L. Pinsky

**Affiliations:** Department of Ecology, Evolution, and Natural Resources, Rutgers University, 14 College Farm Road, New Brunswick, New Jersey 08901, USA; Department of Ecology and Evolutionary Biology, Princeton University, Princeton, New Jersey 08544-1003, USA; Department of Ecology and Evolutionary Biology, University of Toronto, Toronto, Ontario, M5S 3B2, Canada

**Author notes:** Correspondence to: Edward W. Tekwa.

**Keywords:** bifurcation, alternative stable states, *P*-value, coral reef, Lorenz system, conservation behaviour, multiple attractors, regression, finite mixture model

## Abstract

Multiple attractors and alternative stable states are defining features of scientific theories in ecology and evolution, implying that abrupt regime shifts can occur and that outcomes can be hard to reverse. Here we describe a statistical inferential framework that uses independent, noisy observations with low temporal resolution to support or refute multiple attractor process models. The key is using initial conditions to choose among a finite number of expected outcomes using a nonstandard finite mixture methodology. We apply the framework to contemporary issues in social-ecological systems, coral ecosystems, and chaotic systems, showing that incorporating history allows us to statistically infer process models with alternative stable states while minimizing false positives. Further, in the presence of disturbances and oscillations, alternative stable states can help rather than hamper inference. The ability to infer models with alternative stable states across natural systems can help accelerate scientific discoveries, change how we manage ecosystems and societies, and place modern theories on firmer empirical ground.

## Introduction

Alternative stable states and multiple attractors refer to the multiple possible outcomes of a dynamic system that depend on history (Poincaré, 1885; Strogatz, 2015). Such dynamics weave chance and necessity (Monod, 1971; May, 1977) in contrast to traditional Cartesian functional theories (Descartes, 1637; G.W. Leibniz, 1684; Newton, Isaac, 1846) where chance is simply noise that delays an inevitable outcome. Multiple attractors are fundamental components of ecology (May, 1977; Mumby, Hastings, & Edwards, 2007; Scheffer, Hirota, Holmgren, Van Nes, & Chapin, 2012), speciation (Jacob, 1977; Gould & Lewontin, 1979), cell cycles (Song et al., 2010), human behavior (Skiba, 1978; Brock & Hommes, 1997; Tekwa, Fenichel, Levin, & Pinsky, 2019), and other natural sciences, including thermodynamics (Jaeger, 1998), electronics (Roth & Kinney, 2014), health (Ngonghala et al., 2014), weather (Lorenz, 1963), and cosmology (Layzer, 1975). However, multiple attractor systems continue to confound statistical inference, particularly when only noisy and temporally-sparse observations are available. The central question is: how do we translate multiple attractor process models into testable statistical models?

Analyses of alternative stable states have largely focused on tracking single-site natural or experimental systems through time as a bifurcation parameter changes (Scheffer & Carpenter, 2003; Andersen, Carstensen, Hernández-García, & Duarte, 2009; Scheffer et al., 2009; Song et al., 2010), which requires either expensive manipulation or extensive monitoring in hopes of observing a rare transition. A single-site approach ignores the information from replicate sites with the same underlying dynamics. Moreover, by selectively focussing on cases exhibiting rare transitions, we also miss the considerable information provided by sites that have not transitioned between states. These limitations result in low or ambiguous statistical power and is susceptible to a publication bias that ignores negative findings (Ioannidis, 2005).

One approach that harnesses independent data is to test for multiple modes in outcomes (Hartigan & Hartigan, 1985; Song et al., 2010). However, this approach does not directly test whether multiple modes result from intrinsic nonlinear processes rather than multiple modes in external influences. Other approaches reconstruct the empirical stability landscapes (Song et al., 2010; Hirota, Holmgren, Van Nes, & Scheffer, 2011) or detect signals characteristic of bifurcation (Hsieh, Glaser, Lucas, & Sugihara, 2005; Scheffer et al., 2009), but they also do not test process models. Finally, stochastic versions of dynamic models can be constructed to predict the long-run stationary distribution of outcomes (Cobb, Koppstein, & Chen, 1983; Grasman, Maas, & Wagenmakers, 2009), but this approach only compares empirical outcomes to the highest (“Maxwell convention”) or nearest (“delay convention”) predicted mode, which do not account for historical conditions.

The approaches reviewed above are important sources of empirical evidence for alternative stable states. A common limitation of these approaches is that multiple-attractor predictions can inherently explain more variability in the data than single-attractor predictions (depending on how one chooses among the predictions) and thus should be penalized. Yet standard measures of model complexity, such as Akaike Information Criterion (Akaike, 1974), built into these approaches only penalizes the number of parameters and not the multiple predictions that even trivial models can produce. For example, a sine model can produce infinite attractors that are arbitrarily close to any observation. Such a model explains a large proportion of the variance but is intuitively insignificant (see Supporting Information (SI): A Trivial Sine Model).

Here we describe an approach for fitting and testing models with multiple attractors that uses history (or initial condition) to define a single prediction among the multiple attractors, with the goal of minimizing the artificial explanatory power that multiple attractors can add. The approach harnesses time series of independent replicates of the system but only requires two observations per time series. We use empirical and simulation examples to further address how the method avoids false positives, how it uses information from disturbances that can move systems across basins of attractions, and how it can be extended to systems with oscillatory dynamics. The first two examples exhibit the classic ecological concept of alternative stable states, while the latter represents the more general concept of multiple attractors, where attractors do not necessarily converge on fixed points but on cycles.

## Materials and Methods

The first objective of the paper is to translate process models with multiple attractors into testable statistical models. The target data contains multiple independent time series presumably following the same dynamic processes, each with at least two time points and with known bifurcation parameter values if they occur under different conditions that are accounted for by the process model. For the statistical translation, we employ a finite mixture model (McLachlan, Lee, & Rathnayake, 2019), which attributes component identities – basins of attraction – to individual (final) observations based on historical or initial conditions. The attribution of component identities based on history differs from clustering-based methods (Kaufman & Rousseeuw, 2005), which aim to infer identities from the final observations alone. We first provide an intuition for how to use history to choose among multiple attractors, then formalize the finite mixture procedure.

### Intuition

Given a process model with multiple attractors, we aim to define the expected probability density function across these attractors. One aspect of defining this probability is the mode, or the most likely attractor given a time series. The concept of incorporating history in choosing the attractor can be illustrated through a simple pitchfork bifurcation, with a bifurcation parameter *X* and a response variable *Y*. This system has two attractors after a critical value in *X* (Strogatz, 2015) (Figure 1A, SI: Pitchfork Bifurcation). Let us assign *R* to label the different basins of attraction toward the two stable states: *R=1* for the upper basin, and *R=-1* for the lower basin. A particular time-series observation belongs to one basin or another depending on where it was historically (initial conditions), assuming stationarity of parameters. Then, we transform the solution and observations along a new x-axis defined by *RX* (Figure 1B). The model now predicts at most one stable state for a given *RX*. Notice the stringent adherence of the transformation to the model: datapoints 4 and 5 in Figure 1B have crossed the unstable curve between initial and final observations, yet they are classified according to their original basins. This procedure penalizes a model if a perturbation occurs between time points, or if the real system follows dynamic processes that differ from the proposed model (i.e., the real basins are not those hypothesized). If we instead adopted the delay convention of choosing the attractor closest to the final observation, more variance would be explained but this would in fact be an artificial inflation (SI: A Trivial Sine Model). The objective of our method is not to maximize variance explained; rather it is hypothesis testing in order to gain confidence about the model’s validity.

**Figure 1.**
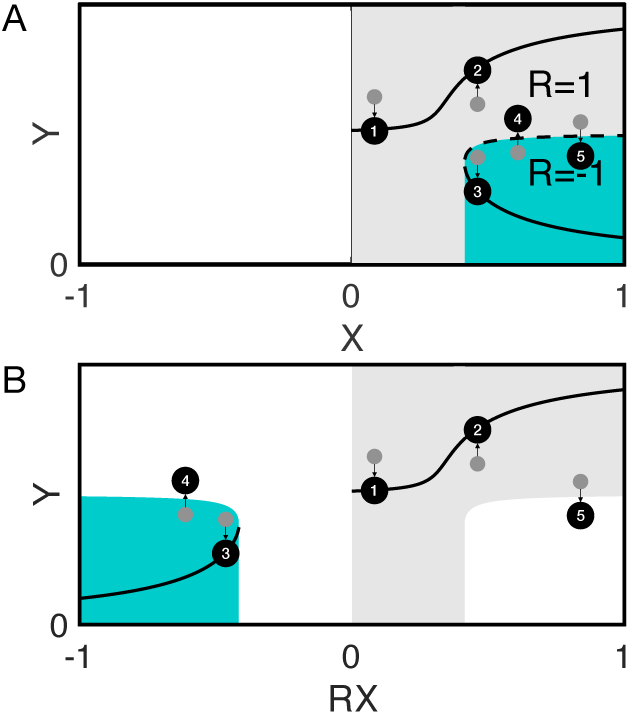
Models with multiple attractors and the empirical utility of history. **(A)** A pitchfork bifurcation features a single solution *Y* when *X* is below a threshold (and above 0), and three solutions above the threshold: two stable (solid curves) and one unstable (dotted curve). The grey and green regions represent the two basins of attraction, which are assigned the values 1 and −1 for the dummy variable *R*. Sample data consist of initial (grey circle) and final (black circle) states. **(B)** Plotting *Y* against *RX*, the transformed bifurcation variable, yields a one-to-one relationship. All data with initial states in the grey region remains in the positive quadrant, while all data with initial states in the green region are reflected onto the negative quadrant.

Even though visualizing *RX* (Figure 1B) becomes difficult when there are more than two attractors, or when *X* is multivariate or includes negative values, all that is required in practice is that for each datum, its initial state chooses which of the alternative attractors lock in. This can be expressed as:

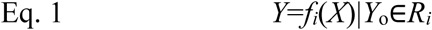

*Y*_*o*_ is the observed initial state, and *Y* is the expected outcome. The function *f*_*i*_(*X*) is specific to the basin of attraction (*R*_*i*_) to which *Yo* belongs. Thus, we establish a way to incorporate history in the relationship between a causal candidate *X* and a single time-invariant outcome *Y*. In technical terms, conditioning prediction on history provides a solution to the implicit regression problem of an irregular type (Hartelman, Maas, & Molenaar, 1998).

### Definition

We now formalize the process and consequences of choosing among attractors in terms of a finite mixture model, which replaces the single outcome expectation *Y* with a distribution of the response variable *y*. Let *g* be the probability density function of *y* at a given predictor value X. The function is:

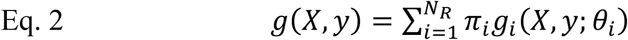

In the equation, *N*_*R*_ is the number of components or basins of attraction, *π*_*i*_ is the mixing proportion of component *i*, and *g*_*i*_ is the component probability density. *θ*_*i*_ specifies the component density parameters, which can include the mode (*Y*_*i*_) and other shape parameters of Gaussian, beta, or other standard distributions depending on application. These parameters correspond to the equilibria in the process model. We employed a nonstandard mixture (Panel on Nonstandard Mixtures of Distributions, 1989) by specifying *π*_*i*_ as a function of initial condition *Y*_*o*_. In particular, one component distribution *g*_*i*_ is chosen (*π*_*i*_=1) based on initial condition *Y*_*o*_ and all other components become degenerate (*π*_*j≠i*_=0). Eq. 3 defines this mixture procedure (following Eq. 1):

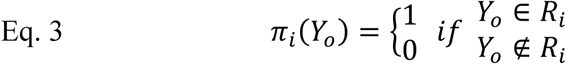

This approach, which assumes parametric component distributions (*g*_*i*_) *a priori*, contrasts from generating the error distribution using the process model (via stochastic differential equations (Grasman et al., 2009)). Our approach attains a simpler and computationally more efficient statistical model from a multiple attractor process model.

### Test Statistics

Finally, armed with probability distributions and thus likelihoods of observations, we can obtain parameter estimates via maximum likelihood, confidence intervals via bootstrapping (B. Efron & R. Tibshirani, 1986), and *P*-values to test the null hypothesis via permutation (Anderson, 2001). Parameter estimates are important for multiple attractor models just like for traditional models when the parameters cannot be directly measured. The probability distribution defined in Eq. 2 and Eq. 3 allows us to first obtain the likelihood (*g*) of an observed final state (*y*) given a parameterized model, then fit the model by iteratively search for the parameters that give the maximum likelihood. The same fitting procedure can be performed repeatedly on bootstrapped data (sampling the *X-*(*Y, Y*_*o*_) sets with replacement) to obtain a distribution of parameter estimates which generates confidence intervals, thereby quantifying uncertainties about components of the process model.

In addition, hypothesis testing can be performed by repeatedly fitting the model to permuted data, with the permutation (sampling without replacement) being performed on the observed initial and final states (*Y, Y*_*o*_) as intact pairs. Alternatively, the predictors (*X*) can be permuted, with the objective being that *Y* and *Y*_*o*_ are decoupled from *X*. The model fits to multiple permuted datasets yield the frequency distribution of test statistics, such as likelihood or variance explained, given that the null hypothesis is true; i.e., given that there is no relationship of the form that the multiple attractor model predicts. By comparing how frequent the null distribution of fits equals or exceeds the original model fit, we obtain the *P*-value, with a low value supporting the multiple attractor model over the null model.

This minimal set of statistics conveys not only how well a model explains the data, but also how likely data with no relationship between *X* and *Y*, *Y*_*o*_ (the null hypothesis) would generate an equal or better fit. This latter question of fit depends on both model complexity (in terms of the number of free parameters) and the number of attractors the model can predict. The key feature of this methodology is that the multiple attractor model’s finite equilibria, further narrowed to one prediction by the initial condition, can easily be wrong. The fit to data is not trivially inflated by multiple predictions or temporal autocorrelation, unlike autoregressive models where lagged states are directly correlated with final states. This set of statistical procedure resembles traditional regressions (A.M. Legendre, 1805) and inherits their advantages (easy to interpret and adaptable to different models) and disadvantages (is affected by oscillatory dynamics). The standard assumptions of independence between time series and identical dynamic processes across series apply; on the other hand, differences in parameter values across series are captured in the bifurcation parameter. The issue of oscillatory dynamics will be addressed through simulations in the Results.

The inferential steps introduced here are demonstrated using three examples of ascending dimensionalities and dynamic possibilities that are drawn from social-ecological systems, coral ecosystems, and chaotic systems. Matlab code and data are available on an online repository: http://github.com/pinskylab/Multiple-Attractor-Inference.

## Results

Here we present three cases of increasing complexity where multiple attractor model inference is performed.

### Empirical Application in a One-Dimensional System

We first study a one-dimensional model that predicts whether human institutions decide to over-harvest or to conserve a renewable resource (Tekwa et al., 2019). The objective here is to demonstrate an inference of alternative stable states using real data from a fishery bioeconomic system. We will demonstrate how to obtain parameter estimation, confidence interval, and hypothesis testing statistics.

Institutions are assumed to adjust their harvest rates in order to maximize the instantaneous economic rent derived from harvesting marine life. We assume that the growth of a biological resource *S* is a logistic function with intrinsic growth *r*, competition *a*, and is reduced due to a harvest rate *F* (the portion of standing stock that is harvested). The economic parameters are the empirically known average cost-to-benefit ratio *γ*, and the unknown number of substitutable stocks (per stock) *N*. The one-dimensional dynamic equation for harvest rate is:

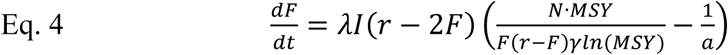

This equation can be solved to obtain the equilibria in *F*. Let the normalized harvest rate solutions *U\** (Eq. 5) be log_2_(*F\**/*F*_*MSY*_), where *F*_*MSY*_=*r*/*2*. The equilibria are:

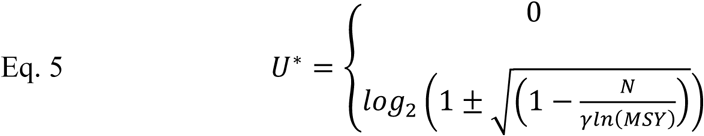

The term *γln*(*MSY*) is a bifurcation parameter, which we label *X*. Stability analysis (Tekwa et al., 2019) shows that when *X<N*, *U*^*^=0 is the only stable equilibrium. When *X>N*, *U*^*^=0 becomes unstable and the two solutions specified by the second line of Eq. 5 become stable.

We obtained empirical assessments of *F*, *F*_*MSY*_, *MSY* and *γ* across 217 marine fisheries (Tekwa et al. 2019). The free parameter *N* was estimated by iteratively searching for the maximum likelihood, which—assuming Gaussian error—is equivalent to maximizing *R*^*2*^. The residuals were computed based on each fishery’s distance to the stable solution (*U*^*^) that the fishery’s initial condition chose, i.e., whether the initial harvest rate *U*_*o*_ was above or below *U\**=0 (according to Eq. 1). Previously, we reported that the pitchfork model explains a moderate portion of variance (*R*^*2*^=0.29) (Tekwa et al., 2019). Here we further show confidence intervals for model predictions, which were obtained from bootstraps by repeatedly (2000 times) refitting the model to 217 random samples with replacement from the original dataset (Figure 2A, blue shade) (B. Efron & R. Tibshirani, 1986). Bootstrapping estimates the sample distribution of the *R-X-Y* relationship and therefore the precision of the parameter estimates. The confidence intervals are not indicative of the observed high data variance, but rather of uncertainty in the parameters of the pitchfork model, which is low in this case.

**Figure 2.**
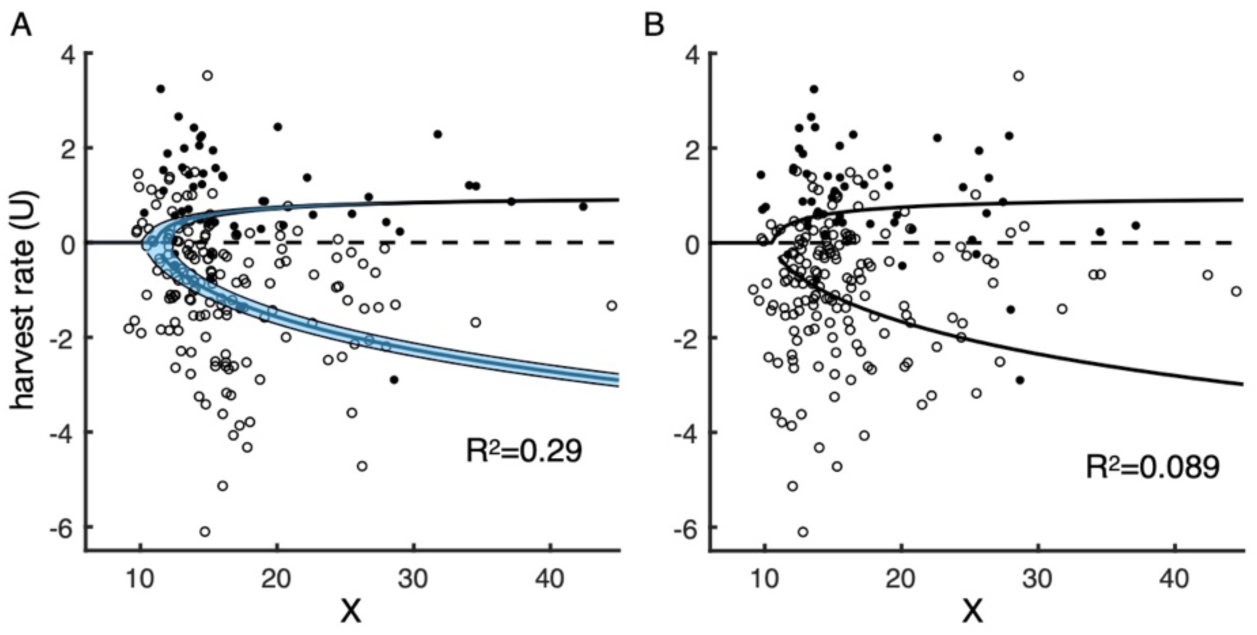
Harvest rate model fit and permutation test. Harvest rate *U*>0 indicates over-harvesting. The bifurcation variable *X* is proportional to a cost-benefit ratio and the maximum sustainable yield of the renewable resource. The bifurcation occurs at *N*, a free parameter that is estimated. **(A)** Model fit to global fishery data (*n*=217), with solid curves indicating stable states (with bootstrapped 95% confidence interval in blue) and a dotted line indicating the unstable state. Filled circles are final observed states whose initial states were above the unstable state, while open circles are final observed states whose initial states were below the unstable state. Estimated *N*=11.2 [10.3-12.2], and *P*=0.7×10^−4^ from the permutation test. **(B)** Model fitted to a dataset after permuting *X* by stock. Such permuted and re-fit estimates contribute to the null *R*^*2*^ distribution used to obtain *P*-values.

A hypothesis test is necessary to provide confidence that a multiple attractor model is not only fitting to the data because it is inherently more flexible. We use a permutation test, with the null hypothesis of no relationship of the modelled form (Eq. 5) between *γ*ln(*MSY*) and *U*. For each permutation set, the harvest rates *U* and *U*_*o*_ were sampled as intact pairs without replacement and were decoupled from *γ*ln(*MSY*). The one-sided *P*-value was obtained from permuting *X* values 10^5^ times and fitting the model to the permuted data to obtain a null distribution of *R*^*2*^ (Anderson, 2001; Tekwa et al., 2019). Given *n*_*P*_, the number of times that the null *R*^*2*^ is greater than the original *R*^*2*^, the *P*-value is *P*=(*nP*+1)/(10^5^+1). We illustrate the process further with a particular instance of permuted data fit (Figure 2B), which yielded *R*^*2*^=0.089, a low but non-zero fit. This fit illustrates how non-linear models with predictions conditional on history could explain some portion of variance even when the data comes from a system without the hypothesized dynamics. Nevertheless, the one-sided *P*-value – the proportion of the permuted fits that were better than the original fit – was low (0.7×10^−4^). This test allows us to conclude it is unlikely that data not exhibiting the hypothesized alternative stable state relationship would result in as good a model fit as observed. Model comparison can also be performed similarly by comparing likelihood-based AIC scores of alternative models, including those without multiple attractors, but these will not be covered here because there are many (infinite) possible alternative models. In a previous paper (Tekwa et al., 2019) we showed that alternative bioeconomic models performed worse in comparison to the alternative stable state model. The results show that multiple attractors can be inferred from real data, with confidence intervals and *P*-values contributing previously unavailable evidences capable of falsifying or supporting multiple-attractor theories.

### Disturbances in a Two-Dimensional System

While empirical inference is the ultimate goal of our methods (e.g., the previous example), here we use simulated data to test the methods where the true data-generating process (and hence the answers to statistical inference) are known in order to reveal subtleties about the method. We examine a two-dimensional coral-macroalgal dynamics model (Mumby et al., 2007), which is similar to other bistable systems that contain a fold or cusp bifurcation (Thom, 1972). Two-dimensional systems contrast with the previous one-dimensional example in that the separatrix—the curve that separates the basins of attraction—is also two-dimensional, generally non-linear, and requires non-equilibrium analysis. In particular, we explicitly explore how disturbances (Hughes et al., 2019), traditionally a source of mismatch between theory and data, provides an opportunity to discover potential basins of multiple attractors. In particular, we quantify fit, perform null hypothesis testing, and compare how an alternative linear model performs as time elapses since disturbance.

The model’s state variables are proportional cover of coral and macroalgae (summing to a maximum of one) (Mumby et al., 2007). The five parameters are coral birth rate (*b*), coral mortality (*d*), macroalgal birth rate (*γ*), macroalgal overgrowth rate on coral (*m*), and grazing rate (*g*). We assume *b=*1 and *m=γ*, which simplifies but retains the full model’s conditional bistability. The dynamic equations are:

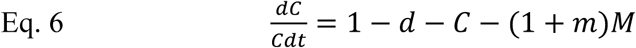

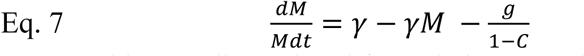

This two-dimensional formulation was obtained from a dimensionality reduction of the original Mumby model (SI: Coral-Macroalgae Model). The grazing rate *g* of fish and invertebrates on macroalgae is a bifurcation parameter such that coral cover has a low equilibrium at low grazing rates, a bistable region at intermediate grazing rates, and a high equilibrium at high grazing rates due to reduced growth of the competing macroalgae (Figure 3A). Unlike the previous one-dimensional system where the basin of attraction can be determined by whether the initial state is above or below the unstable equilibrium (analogous to Figure 3A dashed curve), both initial coral and macroalgal covers must be used to numerically determine the basin of attraction (Figure 3A open and filled circles, SI: Coral-Macroalgae Model). We followed six replicate sets of 40 time-series, each with random initial coral and macroalgal covers that were far from equilibria and thus emulated disturbances at time zero. Final coral covers, for model evaluation purpose, were measured at eight time steps (i.e., years) between *t=*0 and 64. As time elapses from disturbance, systems should converge on the equilibria chosen by the historical disturbances.

**Figure 3.**
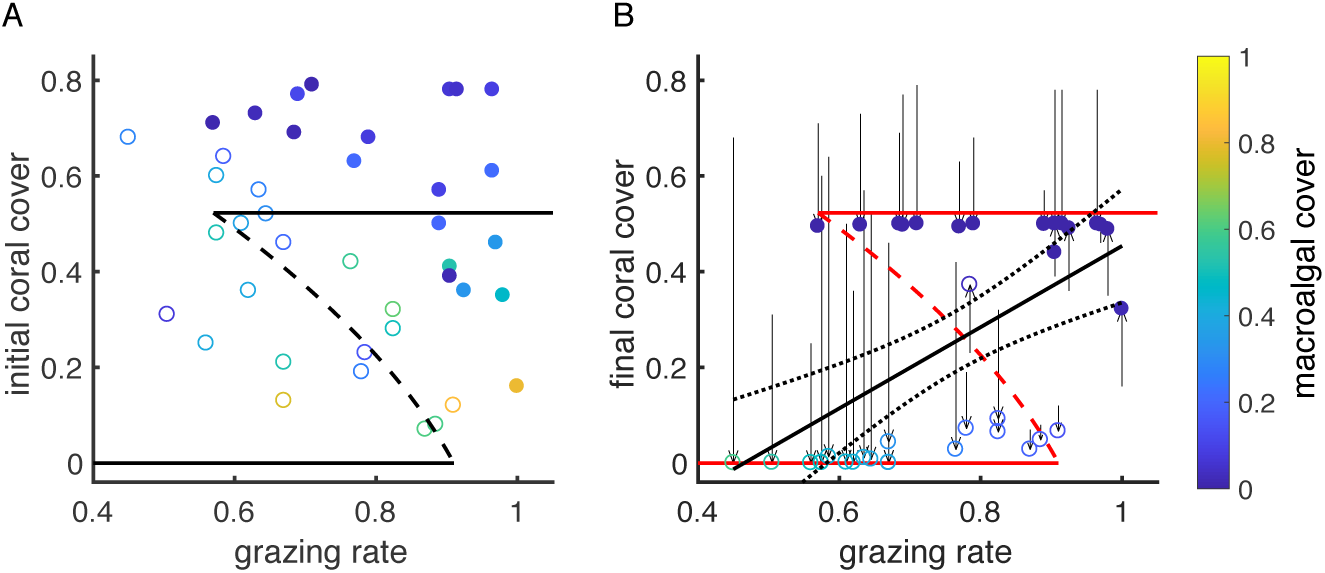
Coral-macroalgal simulated data and model fits. **(A)** Solid (stable) and dashed (unstable) curves represent the true model, and open and filled circles represent random initial states that belong, respectively, to the lower and higher coral cover basins of attraction (*n*=40). Color represents initial macroalgal cover (see color bar). **(B)** The dynamic system was run to t=16, when the final states were labelled as open (lower attractor) or filled (upper attractor) circles depending on the best-fitting model (red curves, *ρ*^*2*^=0.92, *R*^*2*^=0.89). As an alternative model, the black solid and dotted lines are the mean and 95% confidence intervals for a linear regression (*ρ*^*2*^=0.30). Color represents final macroalgal cover (see color bar).

We estimated the two free parameters *d* and *γ*, plus a measure of precision at various times since disturbance for each replicate set through iterative maximum likelihood. Precision is defined by the beta distribution, which we use as the probability density, or the likelihood, of each observation given a predicted mode. The predicted mode depends on *g* and initial conditions. We numerically determined each site’s basin of attraction by running the parameterized differential equations for a sufficiently long time (*t*=64), then choosing the solution closest to the modelled end state. Note that as the parameters vary in search of the best estimates, the predicted basins of attraction shift such that the estimated classification of data by initial condition can be different from their true classification. One instance of a best-fit model is shown in Figure 3B (red).

We observed that with greater time elapsed, the parameter estimates converged on the true values and precision increased (Figure 4A). Model fit was quantified by:

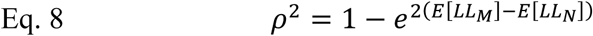

**Figure 4.**
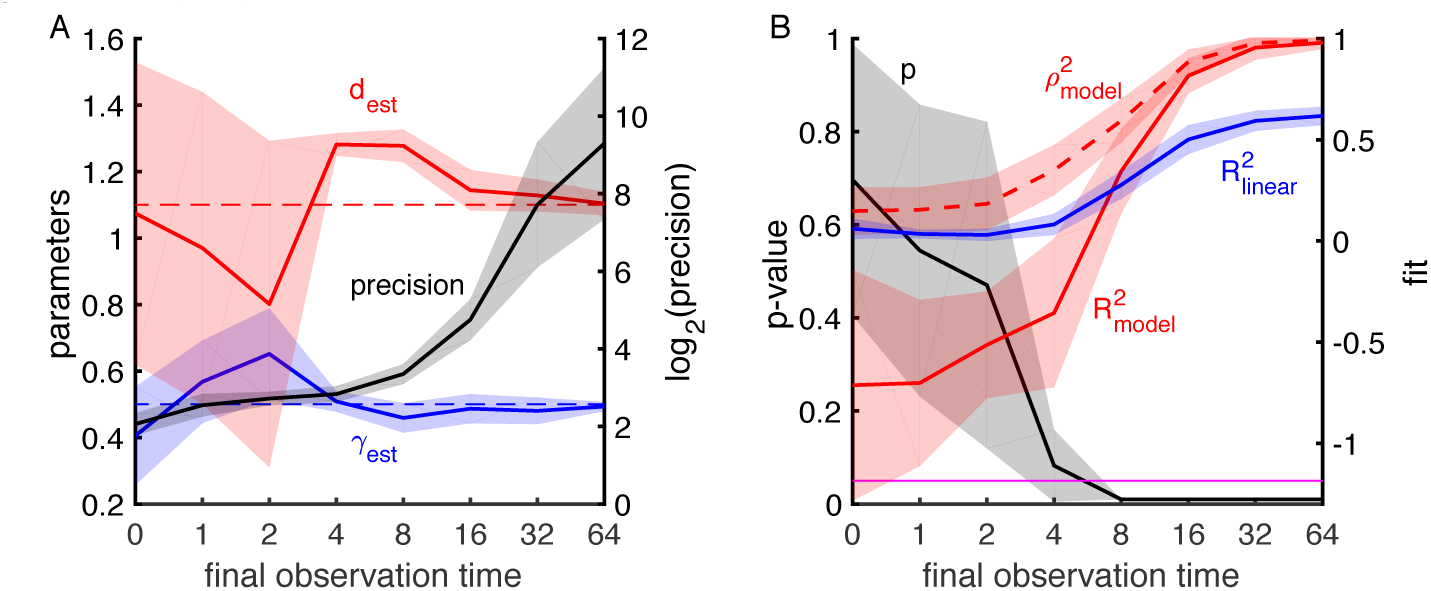
Parameter estimates and model fits for simulated coral-macroalgal covers at different observation times since disturbance. **(A)** Two parameters, coral mortality (*d*_*est*_, red) and macroalgal birth rate (*γ*_*est*_, blue), were estimated for six sets of 40 observations with different initial states and compared to true values (*d*_*true*_, *γ*_*true*_, dashed lines). Precision was also estimated (black line, right axis). **(B)** Fit of a linear regression model was measured as *R*^*2*^ (blue). Fit of the bistable model was measured as *ρ*^*2*^ (beta distribution, red dashed) and *R*^*2*^ (red). *P*-values are computed by permutation tests for the model (black). A magenta horizontal line indicates *P*=0.05 as a reference. Shades are 95% confidence intervals.

*LL*_*M*_ is the log-likelihood of the model and *LL*_*N*_ is the log-likelihood of the mode (most likely value) fitted to the data (Kent, 1983). *ρ*^*2*^=*R*^*2*^ if error is assumed Gaussian; both *ρ*^*2*^ and *R*^*2*^ are plotted in Figure 4B to allow for easier comparison with an alternative (but wrong) model: a linear regression. For both the linear regression and the estimated bistable model, *R*^*2*^ and *ρ*^*2*^ increased with time (Figure 4B, red and blue lines). As might be expected, we found that a linear regression explains the data reasonably well despite the bistability of the underlying dynamics. However, with sufficient time since disturbance, the estimated bistable model eventually explained more variation in the data. We next conduct permutation tests to show how time since disturbance affects hypothesis testing and false positive rates.

The permutation test assessed the *P*-value of the bistable model by asking how often the model fit permuted data (randomized grazing rate *g* 100 times) better than it fit to the original data. The null hypothesis is that there is not a bistable relationship of the form predicted by Eq. 6 and Eq. 7 between *g* and *C*. At time zero when system states were purely random, low *P*-values would indicate false positives. Assuming that *P*=0.05 is a cutoff for a positive result, the false positive rate was 0.038 according to the normal distribution of *p*-values, which was close to the expectation (of 0.05) (Figure 4B, black line). The mean *P*-value dropped below 0.05 between times of 4 and 8 since disturbance, after which time the systems had sufficiently converged on the equilibria to be detectably better than the null model.

We conducted an additional test for false positive rate based on model comparison. We fit both the bistable model and the linear model to simulated noisy linear datasets that correspond to the linear relationship shown in Figure 3B. The bistable model should not fit the data as well as a linear regression. We then compare the models’ likelihoods (Akaike, 1974), which showed the bistable model was a better fit than the correct linear model with a frequency of 0.041 (i.e., the false positive rate, Figure S1, SI: Coral-Macroalgae Model).

These results show that at and right after a disturbance when the response variable is randomly distributed, or when the underlying dynamics are linear, our method has an acceptably low rate of false positives for identifying bistability. In the bistable system, soon after the disturbance when time series begin to converge on equilibria, we were able to infer the correct bistable model. In general, the time lag after disturbance necessary to ensure reliable statistical inference of the underlying dynamics will depend on disturbance magnitude, frequency, and the speed of the dynamics. Initial observations made after large disturbances rather than before should also help in predicting the correct subsequent states.

### Oscillations in a Three-Dimensional System

Finally, we apply our model evaluation method to the three-dimensional Lorenz system (Lorenz, 1963). The system is a classic example of chaos or the butterfly effect in complex systems such as atmospheric convection (SI: Lorenz System) and biological communities (May, 1976; Schaffer, 1985). However, it also features a separate regime exhibiting multiple attractors, whereas most biological models only show chaos, oscillatory behaviour, or the relatively trivial multiple attractors of coexistence and extinction (May & Oster, 1976; Hastings & Powell, 1991; Gakkhar & Naji, 2003; González-Olivares et al., 2011). A tri-trophic model with nonlinear allometries can produce both multiple attractors and chaos (McCann & Yodzis, 1995), but it contains more parameters than the Lorenz system and exhibits similar dynamic possibilities. Here the purpose of using the Lorenz system is not to improve prediction *per se*, but to examine how the coexistence of both multiple attractors and oscillations presents a challenge to our proposed data-sparse method. We demonstrate how multiple attractors assist inference in the presence of oscillation.

The equations for the state variables *X*, *Y*, and *Z* are:

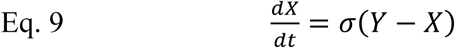

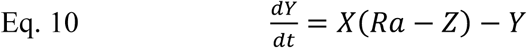

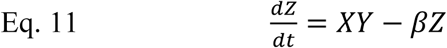

The fixed parameters are *σ*=10, *β=*8/3, and fixed initial conditions of *Y*(0)=1 and *Z*(0)=1. This setup leaves *Ra* as a bifurcation parameter. We illustrate the dynamics of the system using a bifurcation diagram, which shows the points where a trajectory stays (stable points) or turns (extrema in an oscillation). For *Ra* ranging from 1 to 45, in steps of 0.1, we initialized 40 time series with *X*(0) ranging from −60 to 60 in steps of 0.25. Each time series was run for 1000 time steps, and 10 extrema of each series, meaning 10 values of *X* where *dX/dt* were closest to zero (i.e. the points visited), were recorded and plotted in Figure 5A. The expected states of the true model are E[*X*] (across 1000 time points) for each *Ra* and *X*(0). For example, at *Ra*=24, the plot of E[*X*] versus *X*(0) (Figure 5A, top) illustrates the expected state of a time series given an initial *X* value.

**Figure 5.**
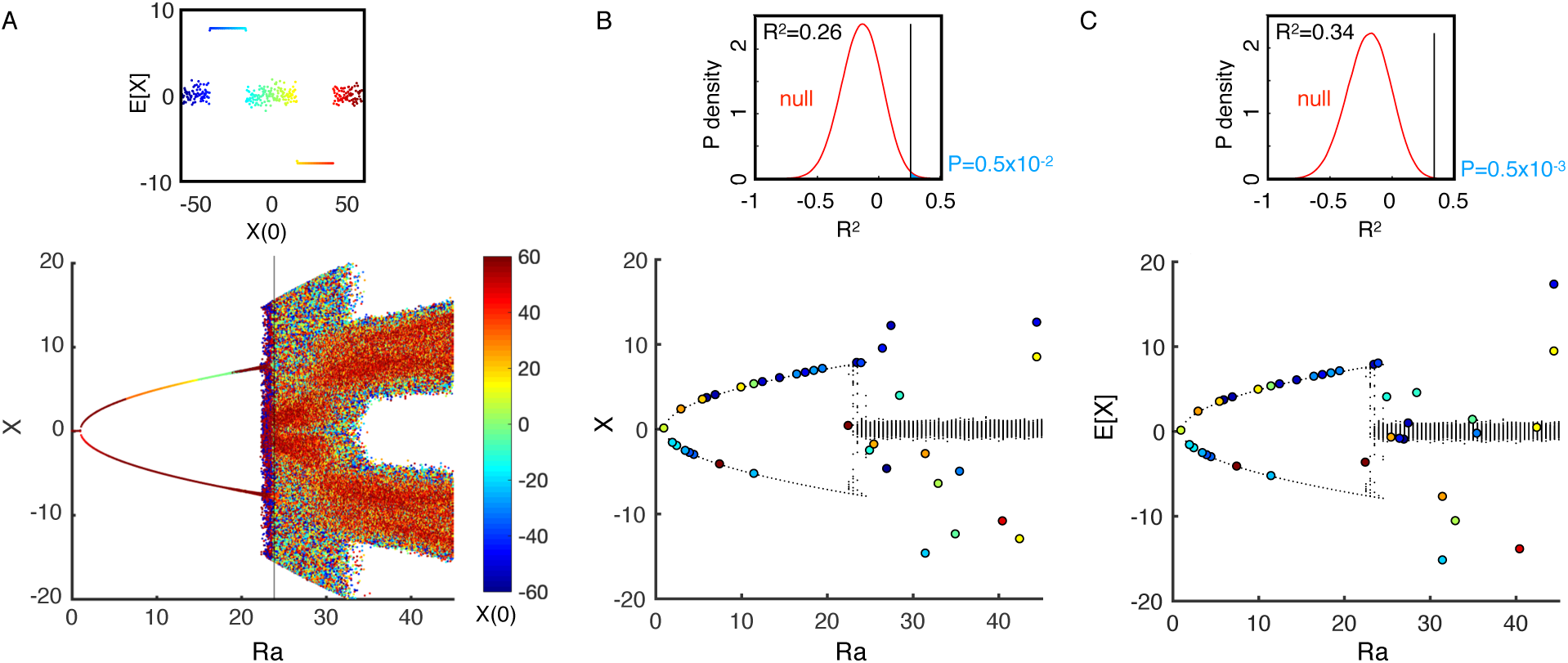
Lorenz system fit to simulated data. **(A)** Extrema (*dX/dt*=0) of the system’s convection rate (*X*), for a range of parameter *Ra* values and initial conditions (*X*(0), color bar). At *Ra* =24 (vertical line), the model is a mix of two stable states and an oscillatory attractor with expected outcome E[*X*] close to 0 (subplot on top). **(B)** The model’s expectations E[*X*^*^] (black dots) are evaluated against 40 simulated observations, with known initial states (color) and final states (position of circles on y-axis) taken at random times for each time series. The simulated observations were permuted to create a null distribution of *R*^*2*^ values (red curve in subplot on top), which was greater than the unpermuted *R*^2^ (vertical line) a portion *P* of the times (blue area). **(C)** The model’s expectations were again evaluated as in **(B)**, but with final states being averages of measurements at two random time points.

For *Ra* <24.7, there are two stable equilibria associated with a pitchfork bifurcation, but they can be accompanied by oscillatory attractors. At *Ra* >28, a single chaotic (strange) attractor with E[*X*]=0 (Lorenz, 1963) emerges, as seen in the high number of extrema (Figure 5A). Observations at low temporal resolution (i.e., at lower than the Nyquist rate (Nyquist, 1928)) cannot reconstruct oscillations and phases, which casts doubt as to whether low-resolution independent time series can be harnessed to infer multiple-attractor models even when there is no process or observation error. We propose that observed states can be compared to a model’s mean long-run predictions (E[*X*^*^]) rather than to single solutions *X*^*^ in the previous cases of stable equilibria. These predictions would still use history and should yield meaningful *P*-values that facilitate inference.

We measure the initial condition and a subsequent state at random times in each time series across *Ra* (Figure 5B circles). We then compute each time series’ residual against the long-run average of the attractor (E[*X*^*^], the predictions, Figure 5B black dots) whose basin of attraction the series’ initial condition belongs. Using residuals, we evaluate *R*^*2*^ and *P*-value from reassessing *R*^*2*^ in datasets where *Ra* is permuted 10^6^ times (Figure 5B subplot). The null hypothesis is that there is not a Lorenz model relationship between *Ra* and *X*. Although the variance explained appeared low (*R*^*2*^=0.26) due to intrinsic oscillations, history-dependence in the bistable region made it unlikely that a null model would produce such a fit (*P*=5×10^−3^).

Inference can be improved under oscillatory dynamics if we can measure initial state and more than one subsequent state within each time series. For example, if the final observation is taken to be the mean of two measurements at different times, *R*^*2*^ increased and *P*-value decreased (Figure 5C). The Lorenz case is challenging because the oscillatory regime essentially gives no information with low temporal measurements (E[*X*^*^]≈0 for *Ra*>24.7), but it illustrates that data in the multiple-attractor regime (low *Ra*) can be instrumental rather than confounding in statistical inference.

## Discussion

We established a statistical fitting and test procedure for bifurcation models that exhibit alternative stable states and multiple attractors. The power of this statistical procedure rests in harnessing observational information from multiple independent time-series from replicates with the same process dynamics, but which requires no more than two time-points within each time series. These requirements align well with the spatially replicated but temporally patchy and short nature of typical ecological time series. We demonstrated model fitting with one and two free parameters in empirical and simulation cases, obtained confidence intervals, and obtained *P*-values for the model against the null hypothesis of no relationship between *X* and *Y*. We also found reliable false positive rates when we compared multiple-attractor and linear model performances against both bistable and linear data. Our focus on model-specific hypothesis-testing approach complements a diversity of existing time-series and observational methods that aim to corroborate alternative stable states (Cobb et al., 1983; Hartigan & Hartigan, 1985; Scheffer & Carpenter, 2003; Hsieh et al., 2005; Andersen et al., 2009; Scheffer et al., 2009; Grasman et al., 2009; Song et al., 2010; Hirota et al., 2011).

A potential problem with incorporating history in prediction, as we do here, is that they are not separable in a causal pathway; i.e., one cannot completely remove the signature of history from predictions since they belong to the same dynamically linked time series (Takens, 1981). The non-separability of history and outcome may be a defining feature of statistical models with multiple attractors, which mix features of autoregressive models – models that use previous states to predict future states – with ordinary regressions. Our approach goes some way to alleviate the problem, in that here the historical state chooses a process-determined solution rather than predicting the final state through autocorrelation as is typical in autoregressive models. The permutation test further addresses this issue by focussing on how much variation in the data the bifurcation variable explains while retaining the original relationship between history and prediction in constructing the null distribution of the test statistic (e.g., likelihood). In other words, we retain in the null distribution the portion of variance explained purely by the influence of initial conditions on the model, thereby reducing the probability of false positives.

The method currently relies on the computational feasibility of iterative maximum likelihood and finding basins of attraction (by running the dynamical equations sufficiently long for convergence). The method therefore shares similar limitations as iterative optimal search methods applied to traditional problems. The method’s ability to identify the true model among competing models will depend on how close the models are to each other and how much data is available. A formal theory of the method’s properties remains to be developed owing to the implicit and irregular nature of multiple attractors (Hartelman et al., 1998), but the method appears to be widely applicable and only limited by computational issues.

Our examples used time series of any length to generate only an initial and a final state, with the goal of utilizing typical ecological data where long series are harder to obtain than site replicates. If long time series are available, our method is data-inefficient, which prioritizes avoiding false positives at the expense of statistical power. In principle however, we can test model fit and other statistics by performing the residual evaluation on every consecutive time point pairs, akin to panel data methods (Arellano & Bond, 1991). This approach could be a venue for future development.

The statistics demonstrated here constitute self-contained, baseline evidence that have long anchored single-attractor theoretical or statistical models (A.M. Legendre, 1805), but are only first steps in scientific inference (Wasserstein, Schirm, & Lazar, 2019). Practical implications and novelty should also be considered as criteria for scientific research in addition to traditional statistics such as *P*-value and confidence intervals (Campbell & Gustafson, 2019). Alternative stable states have strong implications for the management of ecosystems and are of intense intellectual interest (Scheffer, Carpenter, Foley, Folke, & Walker, 2001; Levin et al., 2013). These theories also potentially score highly on novelty criteria such as pre-study probability—the *a priori* probability that a study’s null hypothesis is false, where a low probability indicates novelty (Campbell & Gustafson, 2019). Alternative stable state studies have previously mostly neglected hypothesis-testing statistics and are often select case studies (Schröder, Persson, & De Roos, 2005; Andersen et al., 2009), suggesting that for more inclusive, less curated research with independent replicates, pre-study probabilities may be low and thus highly novel. So far, the combination of novelty and weak evidence has led to the periodic waxing and waning of alternative stable states and multiple-attractor theories throughout the history of ecology and other natural sciences (Monod, 1971; Thom, 1972; Scheffer et al., 2001). The basic ingredients of statistical evidence established here should help improve scientists’ ability to support, reject, and further develop precise dynamic theories.

## Acknowledgements

We thank Simon Levin and Marie-Josée Fortin for discussions. Research was supported by Coral Reef Alliance and The Nature Conservancy PGA-CORAL-042017, NSF awards OCE-1426891 and DEB-1616821, NJ Sea Grant award R/6410-0011, the project Green Growth Based on Marine Resources (GreenMAR, Nordforsk), an NSERC Discovery Grant, and a Canada Research Chair. We declare no conflict of interest.

## Author’s Contributions

EWT, MK, and MLP contributed to the conceptualized the project, EWT conducted the analyses, and all authors contributed to writing.

## Data Availability

Matlab code and data are available on an online repository: http://github.com/pinskylab/Multiple-Attractor-Inference

## References

Akaike, H. (1974). A new look at the statistical model identification. IEEE Transactions on Automatic Control, 19(6), 716–723. doi:10.1109/TAC.1974.1100705

A.M. Legendre. (1805). Nouvelles méthodes pour la détermination des orbites des comètes. Paris: Firmin Didot.

Andersen, T., Carstensen, J., Hernández-García, E., & Duarte, C. M. (2009). Ecological thresholds and regime shifts: approaches to identification. Trends in Ecology & Evolution, 24(1), 49–57. doi:10.1016/j.tree.2008.07.014

Anderson, M. J. (2001). Permutation tests for univariate or multivariate analysis of variance and regression. Canadian Journal of Fisheries and Aquatic Sciences, 58(3), 626–639. doi:10.1139/cjfas-58-3-626

Arellano, M., & Bond, S. (1991). Some Tests of Specification for Panel Data: Monte Carlo Evidence and an Application to Employment Equations. The Review of Economic Studies, 58(2), 277. doi:10.2307/2297968

B. Efron, & R. Tibshirani. (1986). Bootstrap Methods for Standard Errors, Confidence Intervals, and Other Measures of Statistical Accuracy. Statistical Science, 1(1), 54–75.

Brock, W. A., & Hommes, C. H. (1997). A Rational Route to Randomness. Econometrica, 65(5), 1059. doi:10.2307/2171879

Campbell, H., & Gustafson, P. (2019). The World of Research Has Gone Berserk: Modeling the Consequences of Requiring “Greater Statistical Stringency” for Scientific Publication. The American Statistician, 73(sup1), 358–373. doi:10.1080/00031305.2018.1555101

Cobb, L., Koppstein, P., & Chen, N. H. (1983). Estimation and Moment Recursion Relations for Multimodal Distributions of the Exponential Family. Journal of the American Statistical Association, 78(381), 124–130. doi:10.1080/01621459.1983.10477940

Descartes, R. (1637). Discourse on Method, Optics, Geometry, and Meteorology (Rev. ed). Indianapolis, IN: Hackett Pub.

Gakkhar, S., & Naji, R. K. (2003). Order and chaos in predator to prey ratio-dependent food chain. Chaos, Solitons & Fractals, 18(2), 229–239. doi:10.1016/S0960-0779(02)00642-2

González-Olivares, E., Meneses-Alcay, H., González-Yañez, B., Mena-Lorca, J., Rojas-Palma, A., & Ramos-Jiliberto, R. (2011). Multiple stability and uniqueness of the limit cycle in a Gause-type predator-prey model considering the Allee effect on prey. Nonlinear Analysis: Real World Applications, 12(6), 2931–2942. doi:10.1016/j.nonrwa.2011.04.003

Gould, S. J., & Lewontin, R. C. (1979). The spandrels of San Marco and the Panglossian paradigm: a critique of the adaptationist programme. Proceedings of the Royal Society of London. Series B. Biological Sciences, 205(1161), 581–598. doi:10.1098/rspb.1979.0086

Grasman, R. P. P. P., Maas, H. L. J. van der, & Wagenmakers, E.-J. (2009). Fitting the Cusp Catastrophe in *R*: A cusp Package Primer. Journal of Statistical Software, 32(8). doi:10.18637/jss.v032.i08

G.W. Leibniz. (1684). A new method for finding maxima and minima. Acta Eruditorum, 467–473.

Hartelman, P. A. I., Maas, H. L. J., & Molenaar, P. C. M. (1998). Detecting and modelling developmental transitions. British Journal of Developmental Psychology, 16(1), 97–122. doi:10.1111/j.2044-835X.1998.tb00751.x

Hartigan, J. A., & Hartigan, P. M. (1985). The Dip Test of Unimodality. The Annals of Statistics, 13(1), 70–84. doi:10.1214/aos/1176346577

Hastings, A., & Powell, T. (1991). Chaos in a Three-Species Food Chain. Ecology, 72(3), 896–903. doi:10.2307/1940591

Hirota, M., Holmgren, M., Van Nes, E. H., & Scheffer, M. (2011). Global Resilience of Tropical Forest and Savanna to Critical Transitions. Science, 334(6053), 232–235. doi:10.1126/science.1210657

Hsieh, C., Glaser, S. M., Lucas, A. J., & Sugihara, G. (2005). Distinguishing random environmental fluctuations from ecological catastrophes for the North Pacific Ocean. Nature, 435(7040), 336–340. doi:10.1038/nature03553

Hughes, T. P., Kerry, J. T., Connolly, S. R., Baird, A. H., Eakin, C. M., Heron, S. F., … Torda, G. (2019). Ecological memory modifies the cumulative impact of recurrent climate extremes. Nature Climate Change, 9(1), 40–43. doi:10.1038/s41558-018-0351-2

Ioannidis, J. P. A. (2005). Why Most Published Research Findings Are False. PLoS Medicine, 2(8), e124. doi:10.1371/journal.pmed.0020124

Jacob, F. (1977). Evolution and tinkering. Science, 196(4295), 1161–1166. doi:10.1126/science.860134

Jaeger, G. (1998). The Ehrenfest Classification of Phase Transitions: Introduction and Evolution. Archive for History of Exact Sciences, 53(1), 51–81. doi:10.1007/s004070050021

Kaufman, L., & Rousseeuw, P. J. (2005). Finding groups in data: an introduction to cluster analysis. Hoboken, N.J: Wiley.

Kent, J. T. (1983). Information gain and a general measure of correlation. Biometrika, 70(1), 163–173.

Layzer, D. (1975). The arrow of time. Scientific American, 233, 56–69.

Levin, S., Xepapadeas, T., Crépin, A.-S., Norberg, J., de Zeeuw, A., Folke, C., … Walker, B. (2013). Social-ecological systems as complex adaptive systems: modeling and policy implications. Environment and Development Economics, 18(02), 111–132. doi:10.1017/S1355770X12000460

Lorenz, E. N. (1963). Deterministic Nonperiodic Flow. Journal of the Atmospheric Sciences, 20(2), 130–141. doi:10.1175/1520-0469(1963)020<0130:DNF>2.0.CO;2

May, R. M. (1976). Simple mathematical models with very complicated dynamics. Nature, 261(5560), 459–467. doi:10.1038/261459a0

May, R. M. (1977). Thresholds and breakpoints in ecosystems with a multiplicity of stable states. Nature, 269(5628), 471–477. doi:10.1038/269471a0

May, R. M., & Oster, G. F. (1976). Bifurcations and Dynamic Complexity in Simple Ecological Models. The American Naturalist, 110(974), 573–599. doi:10.1086/283092

McCann, K., & Yodzis, P. (1995). Bifurcation Structure of a Three-Species Food-Chain Model. Theoretical Population Biology, 48(2), 93–125. doi:10.1006/tpbi.1995.1023

McLachlan, G. J., Lee, S. X., & Rathnayake, S. I. (2019). Finite Mixture Models. Annual Review of Statistics and Its Application, 6, 355–378.

Monod, J. (1971). Chance and necessity: an essay on the natural philosophy of modern biology (1st American ed.). New York: Knopf.

Mumby, P. J., Hastings, A., & Edwards, H. J. (2007). Thresholds and the resilience of Caribbean coral reefs. Nature, 450(7166), 98–101. doi:10.1038/nature06252

Newton, Isaac. (1846). Principia: the mathematical principles of natural philosophy. New York: Daniel Adee.

Ngonghala, C. N., Pluciński, M. M., Murray, M. B., Farmer, P. E., Barrett, C. B., Keenan, D. C., & Bonds, M. H. (2014). Poverty, Disease, and the Ecology of Complex Systems. PLoS Biology, 12(4), e1001827. doi:10.1371/journal.pbio.1001827

Nyquist, H. (1928). Certain Topics in Telegraph Transmission Theory. Transactions of the American Institute of Electrical Engineers, 47(2), 617–644. doi:10.1109/T-AIEE.1928.5055024

Panel on Nonstandard Mixtures of Distributions. (1989). Statistical Models and Analysis in Auditing. Statistical Science, 4(1), 2–33. doi:10.1214/ss/1177012665

Poincaré, H. (1885). L’Équilibre d’une masse fluide animée d’un mouvement de rotation. Acta Mathematica, 7(1), 259–380.

Roth, C. H., & Kinney, L. L. (2014). Fundamentals of logic design (7th ed). Stamford, CT: Cengage Learning.

Schaffer, W. M. (1985). Order and Chaos in Ecological Systems. Ecology, 66(1), 93–106. doi:10.2307/1941309

Scheffer, M., Bascompte, J., Brock, W. A., Brovkin, V., Carpenter, S. R., Dakos, V., … Sugihara, G. (2009). Early-warning signals for critical transitions. Nature, 461(7260), 53–59. doi:10.1038/nature08227

Scheffer, M., Carpenter, S., Foley, J. A., Folke, C., & Walker, B. (2001). Catastrophic shifts in ecosystems. Nature, 413(6856), 591–596. doi:10.1038/35098000

Scheffer, M., & Carpenter, S. R. (2003). Catastrophic regime shifts in ecosystems: linking theory to observation. Trends in Ecology & Evolution, 18(12), 648–656. doi:10.1016/j.tree.2003.09.002

Scheffer, M., Hirota, M., Holmgren, M., Van Nes, E. H., & Chapin, F. S. (2012). Thresholds for boreal biome transitions. Proceedings of the National Academy of Sciences, 109(52), 21384–21389. doi:10.1073/pnas.1219844110

Schröder, A., Persson, L., & De Roos, A. M. (2005). Direct experimental evidence for alternative stable states: a review. Oikos, 110(1), 3–19. doi:10.1111/j.0030-1299.2005.13962.x

Skiba, A. K. (1978). Optimal Growth with a Convex-Concave Production Function. Econometrica, 46(3), 527. doi:10.2307/1914229

Song, C., Phenix, H., Abedi, V., Scott, M., Ingalls, B. P., Kærn, M., & Perkins, T. J. (2010). Estimating the Stochastic Bifurcation Structure of Cellular Networks. PLoS Computational Biology, 6(3), e1000699. doi:10.1371/journal.pcbi.1000699

Strogatz, S. H. (2015). Nonlinear dynamics and chaos: with applications to physics, biology, chemistry, and engineering (Second edition). Boulder, CO: Westview Press, a member of the Perseus Books Group.

Takens, F. (1981). Detecting strange attractors in turbulence. In D. Rand & L.-S. Young (Eds.), Dynamical Systems and Turbulence, Warwick 1980 (Vol. 898, pp. 366–381). Berlin, Heidelberg: Springer Berlin Heidelberg. doi:10.1007/BFb0091924

Tekwa, E. W., Fenichel, E. P., Levin, S. A., & Pinsky, M. L. (2019). Path-dependent institutions drive alternative stable states in conservation. Proceedings of the National Academy of Sciences, 116(2), 689–694. doi:10.1073/pnas.1806852116

Thom, R. (1972). Stabilité Structurelle et Morphogénèse, Essai d’une Théorie Générale des Modèles. New York: Benjamin.

Wasserstein, R. L., Schirm, A. L., & Lazar, N. A. (2019). Moving to a World Beyond “ *p* < 0.05”. The American Statistician, 73(sup1), 1–19. doi:10.1080/00031305.2019.1583913

