## Supplementary material for "Inferring Models with Alternative Stable States from Independent Observations": SI

**Supporting Information for**
**Inferring Models with Alternative Stable States from Independent Observations**

Edward W. Tekwa[1,2*], Martin Krkošek[3], Malin L. Pinsky[1]

See main text for references.

#### *1. A Trivial Sine Model*

We use the following model to illustrate how multiple predictions can inflate apparent explanatory power. Consider a sine function that represents the state dynamics $dY/dt=\sin(Y/\varepsilon)$ where $\varepsilon$ is a fixed parameter. The solutions of $Y$ are $Z\varepsilon\pi$ where $Z$ is any integer. The second derivative $d^2y/dt^2=\cos(Y/\varepsilon)$ is negative when $Y=(2Z+1)\varepsilon\pi$ (odd multiples of $\varepsilon\pi$), which are the stable solutions. Conversely, $d^2Y/dt^2=\cos(y/\varepsilon)$ is positive when $y=2Z\varepsilon\pi$ (even multiples of $\varepsilon\pi$), which are the unstable solutions. For any given bifurcation parameter value $X$, there are multiple (infinite) stable solutions divided by equally spaced unstable solutions. Thus, any observation in $Y$ is within a distance $\varepsilon\pi/2$ away from the closest stable solution. A scheme that uses the distance from an observation to the nearest stable solution as the residual (Grasman et al., 2009) would deduce that the trivial sine model, with $\varepsilon$ fixed at a very small value, can explain close to all variance of any data. On the other hand, a scheme that uses the most likely mode as the stable solution (Grasman et al., 2009) would be indeterminate, since all modes are equal in magnitude in the sine model. This example illustrates that distance from any stable solution or distance from the largest stable solution are both unsatisfactory.

#### *2. Pitchfork Bifurcation*

We use one of the simplest models with multiple attractors to show how history can be used to determine final states. Consider the equation:

Eq. S1 $$\frac{dY}{dt} = Y(X - Y^2) + \delta$$

The solutions ($Y$) form a pitchfork bifurcation, with the bias $\delta$ causing asymmetry in the two stable equilibria (Figure 1A). For the case of $\delta=0$, stability can be obtained from checking whether all eigenvalues of the Jacobian matrix evaluated at a solution are negative. The results are that, in the bistable regime, the upper and lower solutions are stable while the intermediate solution is unstable. For the case of $\delta\neq0$, the solutions can only be found numerically, but it is well known that the resulting solutions have analogous stability properties to the case of $\delta=0$ (Strogatz, 2015).

#### *3. Harvest Rate Model*

We assume that the growth of a biological resource $S$ is a function of intrinsic growth $r$, competition $a$, and the resource mortality due to harvest $F$ (the portion of standing stock that is harvested):

Eq. S2 $$\frac{dS}{dt} = S(r - aS - F)$$

The social utility, or economic rent that an institution derives from harvesting a quickly equilibrating stock $S^*$ (by solving Eq. S2) at harvest mortality $F$ is the sum of a benefit that diminishes with harvest volume ($FS^*$) and a cost that increases linearly with harvest volume:

Eq. S3 $$u = Vln(FS^*) - IFS^*$$

$V$ is a reference marginal benefit, and $I$ is the marginal cost of harvesting a kilogram.

The evolution of harvest mortality $dF/dt$ is assumed proportional to the change in social utility (or rent) with respect to a change in harvest mortality ($\partial u/\partial F$):

Eq. S4 $$\frac{dF}{dt} = \lambda \frac{r-2F}{a}\left(\frac{Va}{F(r-F)} - I\right)$$

$\lambda$ is a speed of change constant $\lambda$. $I$ and $V$ are not directly measurable from single stock data, because some stocks may be substitutable for each other and thus contribute to diminishing returns together rather than as individual stocks. Given an average cost-to-benefit ratio $\gamma$ and the number of substitutable stocks per stock $N$, each of which on average yields $MSY=r^2/(4a)$, we can solve for $I/V$ (Tekwa et al., 2019):

Eq. S5 $$\frac{1}{MSY\ I/V} = \frac{N}{\gamma ln(MSY)}$$

By substituting Eq. S5 into Eq. S4, we obtain:

Eq. S6 $$\frac{dF}{dt} = \lambda I \frac{r-2F}{a}\left(\frac{aV/I}{F(r-F)} - 1\right) = \lambda I \frac{r-2F}{a}\left(\frac{Nr^2}{4F(r-F)\gamma ln(MSY)} - 1\right)$$

This leads to the dynamic equation in Eq. 3.

### *4. Coral-Macroalgae Model*

The original Mumby et al. model (Mumby et al., 2007) was written with three state variables, including coral cover ($C$), macroalgal cover ($M$), and turf cover ($T$):

Eq. S7 $$\frac{dM}{Mdt} = aC - \frac{g}{M+T} + \gamma T$$

Eq. S8 $$\frac{dC}{Cdt} = rT - d - aM$$

Eq. S9 $$T = 1 - M - C$$

However, turf simply acts as empty space. By replacing $T$ in Eq. S7 and Eq. S8 with Eq. S9 and rearranging, we obtain Eq. 4 and Eq. 5.

The beta distribution, with shape parameters $\sigma$ and $\tau$, is appropriate for coral covers because they are capped between 0 and 1. We truncate observations to be between 0.001 and 0.999 so that edge probability densities are non-zero, then match the model prediction at each grazing rate to the beta mode. Precision ($\sigma+\tau$) is the inverse error estimate, which is constrained to be higher than 2 (to ensure finite probability densities) and is high when error is low. The results of fitting with a beta distribution is similar to those obtained from using a normal distribution (i.e. by minimizing least squares of residuals), but the latter assumption is less realistic because it implies that negative or greater than full coral cover (>1) is possible.

A test of false positive rate based on model comparison was conducted by simulating noisy data that follows a linear trend that resembles outcomes from the bistable coral-macroalgal dynamics. The trend is obtained from the linear regression in Figure 3B. Using standard deviation of 0.1 for both intercept and slope, we generated 1000 sets of 40 time series. For each time series, the initial state is two times the final deviation from the true trend, representing systems that converge on the equilibrium through time. The bistable model (assuming beta distribution) and the linear model (assuming normal distribution) were fitted to the datasets. One instance is shown in Figure S1. The rate at which the bistable model's likelihood is greater than the linear model's likelihood, evaluated using normal distribution for both to ensure comparability, is the false positive rate. This test is equivalent to an AIC model comparison (Akaike, 1974) because both models have the same number of free parameters (two). The bistable model's parameter estimates are $d$=0.67±0.056 and $\gamma$=0.90±0.11, with $R^2$=0.28±18. The linear regression's intercept and slope are -0.35±0.087 and 0.80±0.12, with $R^2$=0.51±0.095.

#### ***5. Lorenz System***

The Lorenz system is used as a generic model to explore complex dynamics, but it was originally derived after atmospheric convection (Lorenz, 1963). The state variables originally stood for the rate of convection ($X$), the horizontal temperature variation ($Y$), and the vertical temperature variation ($Z$). The parameters were the Prandtl number ($\sigma$), the physical constant ($\beta$), and the Rayleigh number ($Ra$), with increasing value indicating a transition from laminar to turbulent flow.

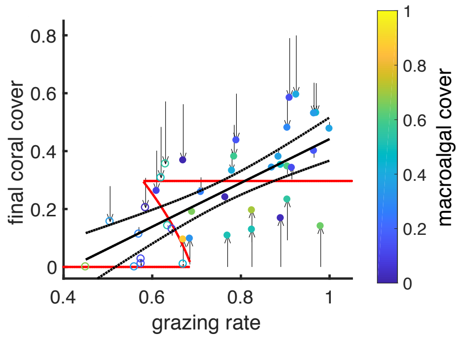


**Figure S1.** Simulated linear data and model fits to test for false positive rate. The 40 final states (dots) were generated randomly around the line defined by an intercept of -0.39 and a slope of 0.85, with a standard deviation of 0.1 for both intercept and slope. Initial states (arrow origins) were recorded as two times the final deviation from the true trend. The final states were labelled as open (lower attractor) or filled (upper attractor) circles according to the best-fitting bistable model (red curves, $R^2$=0.18). The correct model (black solid line with 95% confidence intervals) is a linear regression ($R^2$=0.47). Color represents final macroalgal cover (see color bar). The noisy data and fit comparisons were generated 1000 times to obtain a false positive rate.
